# Calcite Precipitation by a Nitrogen-Fixing Cyanobacterium

**DOI:** 10.1101/2025.02.17.638518

**Authors:** Christian M. Brininger, Jian Wei Tay, Evan B. Johnson, Erin Espelie, Jeffrey C. Cameron

## Abstract

Microbiologically induced calcium carbonate precipitation (MICP) is the process through which the metabolic activity of microorganisms causes the precipitation of calcium carbonate, which can result in solidification of sediment. In cyanobacteria, MICP is thought to occur primarily because cells sequester bicarbonate for the photosynthetic process, thereby lowering the pH of the surrounding media. However, these mechanisms are still poorly understood. Here we show direct evidence of MICP caused by the filamentous cyanobacterium *Anabaena*. *Anabaena* differentiates into photosynthetic vegetative cells and nitrogen-fixing heterocysts. Using quantitative microscopy, we show that MICP occurs due to two distinct mechanisms: Firstly, mechanical stress on vegetative cells can cause leakage and/or lysis, releasing sequestered bicarbonate into the environment, resulting in formation of new crystals. Secondly, contact between a heterocyst and a calcite crystal seed appears to cause rapid crystal growth. Our results suggest an evolutionary benefit of contact-mediated precipitation to anchor cyanobacteria growing in tidal regions. By providing greater insight into MICP caused by *Anabaena*, these results could be used to optimize bio-cement production, thereby enabling a green construction material that could assist with carbon sequestration and reducing the impact of climate change.

## Introduction

Organisms across all kingdoms of life have been shown to influence minerals, such as calcium carbonate (CaCO_3_), as part of their growth. In eukaryotes, organisms such as shellfish utilize calcium carbonate present in the aquatic environment to form vital structures^1,2^. Microorganisms, such as fungi, archaea and bacteria, have also been shown to cause the precipitation of calcium carbonate through different mechanisms, collectively known as microbiologically induced calcium carbonate precipitation (MICP)^1^. Bacteria in particular have been shown to cause MICP through a range of metabolic processes, including urea hydrolysis^3^, denitrification^4^, methane oxidation^5^, sulfate reduction^6^, iron reduction^7^, degradation of calcium oxalate^8^, degradation of amino acids^9^, and photosynthesis^10^.

In recent years, MICP has received attention as a potential tool to address many engineering and environmental issues^11^, for instance to stabilize soil^12^ or as an eco-friendly alternative to traditional cement^13,14^. These materials rely on the formation of CaCO_3_ crystals to both solidify and to provide compressive strength. More recently, cyanobacteria have been shown to be useful as living building materials^15^. Compared to other microbes, cyanobacteria can derive its own energy from sunlight, as well as absorb and store CO_2_ through biological processes. However, MICP mechanisms in cyanobacteria are still poorly understood.

In the ocean and other aquatic environments, cyanobacteria have been observed to influence the precipitation of calcium carbonate during blooms or mass lysis events^16,17^. Several different theories have been proposed to explain this phenomenon. A leading theory is that photosynthesis by cyanobacteria leads to calcium carbonate precipitation by consuming carbon dioxide in the local environment^16^. This in turn drives conversion of bicarbonate ions to reestablish equilibrium: HCO^-^_3_ →CO_2_ +OH^-^. The release of OH^-^ ions increases the pH of the surrounding medium which then drives the precipitation of calcium carbonate, described by the equation^18,19^: Ca^2+^+HCO^-^_3_ →CaCO_3_ +H^+^. However, other studies have shown that calcium carbonate precipitation occurs even in the absence or inhibition of photosynthesis^20^ and theorize instead that adsorption of Ca^2+^ ions by cell-surface proteins are responsible for crystal nucleation^21,22^.

Another popular theory is that cyanobacteria can cause CaCO_3_ precipitation if the cells lyse, for instance through viral infection^23,17^. Cyanobacteria naturally sequester dissolved inorganic carbon (dissolved carbon dioxide when reacted with water) within their cells in order to drive the activity of the enzyme ribulose-1,5-bisphosphate carboxylase/oxygenase (hereafter RuBisCO). Cell rupture leads to the release of bicarbonate ions into the environment, which drives equilibrium once again towards calcium carbonate precipitation.

While all these theories have merit, it is clear that the mechanism(s) of MICP by cyanobacteria is still under debate. To shed light on this matter, here, we utilize time-lapse and multispectral microscopy to film the filamentous cyanobacterium *Anabaena* sp. ATCC 33047 (hereafter *Anabaena*) grown under environmental conditions favoring MICP. This species has been studied recently as a potential candidate for MICP-based engineering applications due to its ability to fix nitrogen, potentially minimizing contaminants during manufacturing^24^. Thus, it is essential to gain a better understanding of how MICP is caused by this species to optimize production yields.

*Anabaena* filaments are typically composed of two main cell types: vegetative cells, which carry out photosynthesis, and heterocysts, which fix nitrogen. By using time-lapse microscopy, we observed unique calcium carbonate precipitation mechanisms caused separately by vegetative cells and heterocysts. Compared to traditional biochemical assays, this approach allowed us to directly track individual cells, as well as crystal formation and growth, thereby providing greater insight into the mechanisms of cyanobacterial MICP.

## Results

### Time-lapse fluorescence microscopy enables individual cells to be tracked

To demonstrate the feasibility of our approach, *Anabaena* was first grown in BG11 media without supplemental nitrogen (denoted BG11_-N_) and filmed under typical laboratory culturing conditions. Cells were grown under a soft agar pad, which constrains cell growth to a 2-dimensional plane. Fig. 1a-c and Supplementary Movie S1 shows representative frames of an *Anabaena* filament. Brightfield and chlorophyll fluorescence images were recorded (see Methods). As the filament grows, some cells differentiate into heterocysts (indicated by carets), which can be identified both by their size and the loss of chlorophyll fluorescence over time.

**Fig. 1.**
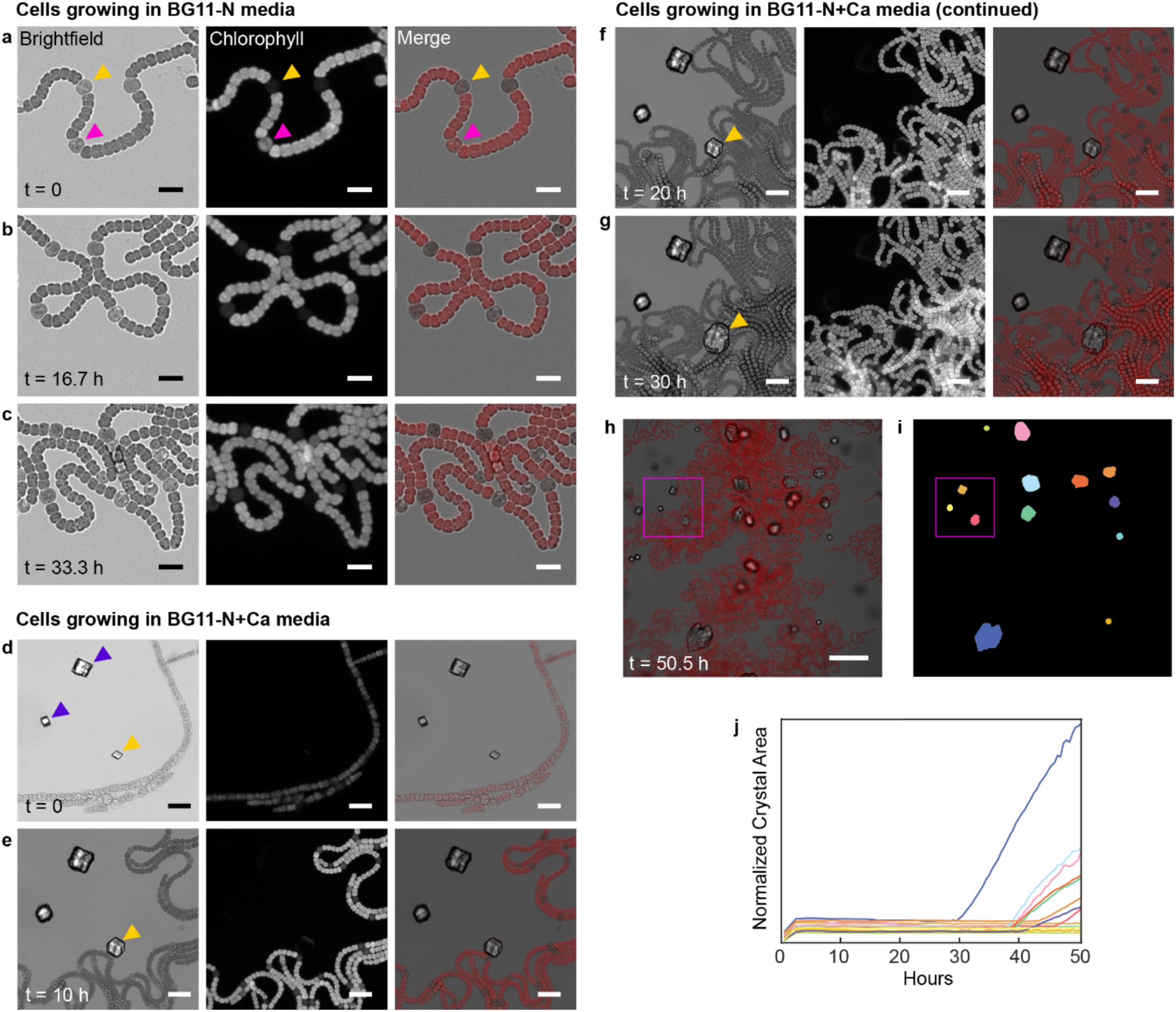
Single-cell analysis of *Anabaena* sp. ATCC 33047 growth and differentiation. **a-c** Representative frames showing Anabaena growing on the microscope in BG-11 media without supplemental nitrogen and without buffer (BG11_-N_). Images were collected in the brightfield and chlorophyll fluorescence channels. A merged image is also shown (chlorophyll fluorescence shown in red). The carets indicate heterocysts, which are identified both by their increased size and the loss of chlorophyll fluorescence. The magenta caret indicates a cell in the early stages of differentiation, and the yellow caret indicates a previously differentiated heterocyst. Scale bars are 10 µm. **d-g** Representative frames showing cells growing in BG11_-N-Buffer_ media supplemented with calcium chloride and bicarbonate (BG11_-N-Buffer_+Ca). Initial seed crystals were observed (indicated by carets). We observed initial crystal growth (seen in frame e), likely due to temperature change as the sample was placed in the growth chamber. Some crystals (yellow caret) showed increased growth due to interaction with the cells. Scale bars are 20 µm. **h** and **i** Crystal growth was tracked over time (Full timelapse shown in Supplementary Movie S3). h shows the merged frame after 50.5 hours and i shows the mask labeling the crystals. **j** Crystal area over time. Scale bars are 50 µm. The zoomed-in region shown in d-g is indicated in magenta.

To observe calcium carbonate crystal growth, 25 mM of calcium chloride and sodium bicarbonate was added to a modified BG11 media without a buffering solution (denoted BG11_-N-Buffer_+Ca). Frames from a representative movie (Supplementary Movie S2) are shown in Fig. 1d-g. We note that this concentration of calcium chloride and sodium bicarbonate caused small seed crystals to form at the start of the movie. We then filmed the cells over time and utilized computational software to track individual crystal growth, as shown in Fig. 1h-j.

As shown in Fig. 1j, we found that all initial seed crystals grew slightly in the first two hours of the movie, likely due to the temperature change when the room temperature pad was placed in the 37 °C growth chamber. Similar growth was observed in experimental replicates. After this initial growth, the crystal area stayed constant for approximately 20 hours. Since the cells continued to grow during this time, our data suggests that photosynthesis alone does not cause crystal growth, at least under these conditions. After this time, some crystals showed growth upon interaction with cyanobacterial cells, as is shown in the following sections.

### Cyanobacterial cell lysis or leakage results in crystal growth

We observed two distinct conditions in which crystal growth occurs. First, cyanobacteria cells can lyse if mechanically trapped against a hard object. Fig. 2a and Supplementary Movie S4 shows an instance in which a vegetative cell grows adjacent to a seed crystal. As the cell grows, one of its progenies becomes confined by its siblings and eventually lyses, likely due to contact against the rough surface of the crystal. This causes measurable crystal precipitation approximately 60 minutes later. Shortly before cell lysis and crystal growth, an increase in chlorophyll fluorescence is observed, which is consistent with our previous observations of the effect of mechanical confinement on individual cells^25^.

**Fig. 2.**
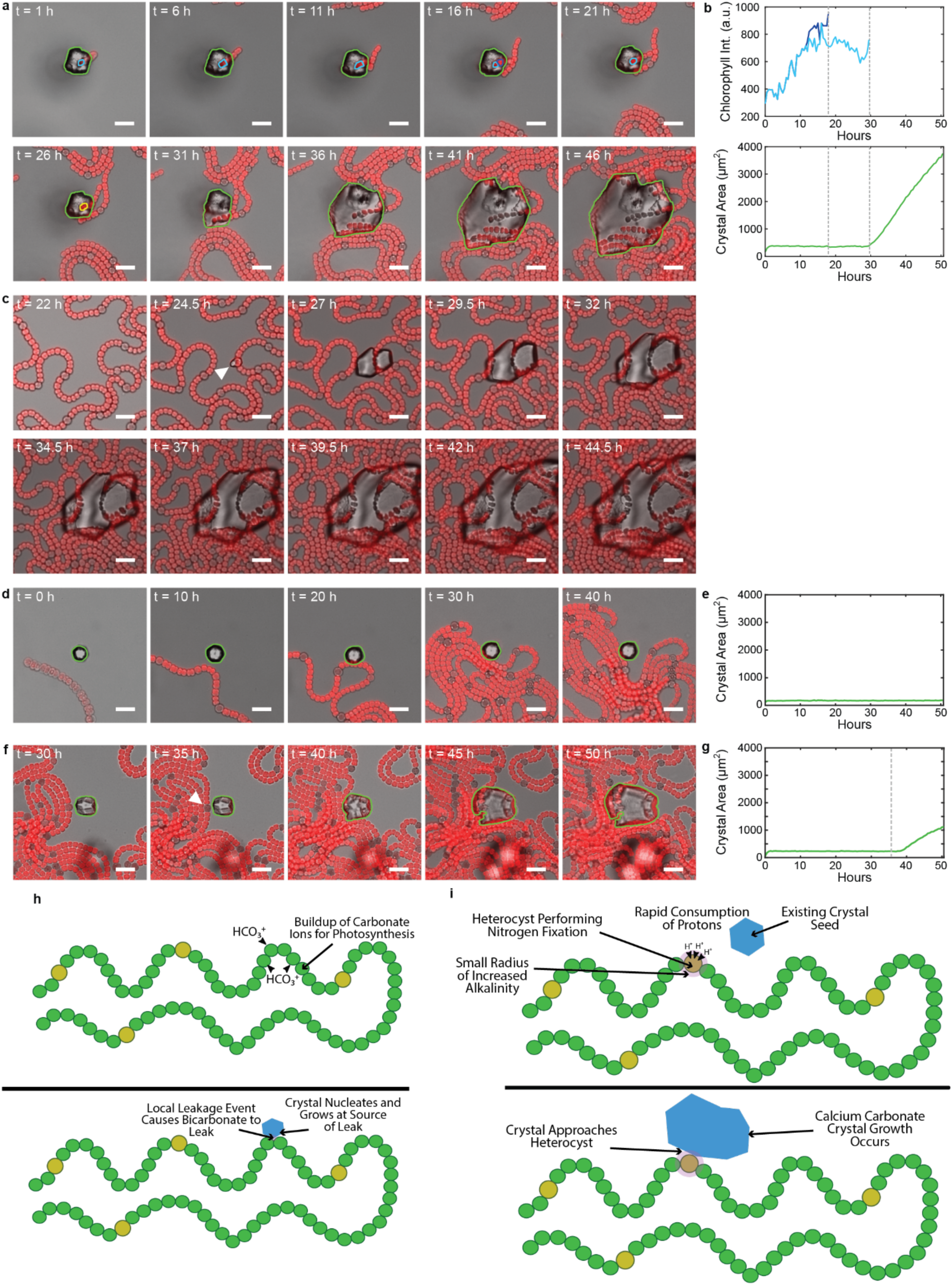
Physical contact with *Anabaena* cells causes crystals to grow. **a** Time-lapse images showing crystal growth as a result of *Anabaena* 33047 cell lysis within a crystal. **b** The average chlorophyll intensity of the two cells tracked and the corresponding growth of crystal area. Dotted lines indicate the time when the individual cells lysed. **c** Time-lapse images showing nucleation and growth of a crystal at a sharp bend of a filament. The crystal growth appeared to be caused by cellular leakage due to mechanical stress at the bend. **d** and **e** Representative frames and plot showing that crystal growth does not occur due to proximity or contact with a vegetative cell. **f** Representative frames showing growth of a crystal after contact with a heterocyst (indicated by an caret). **g** Plot showing crystal area over time. Dotted line indicates time of heterocyst contact at 37 hours. All plots show the brightfield image in grayscale and chlorophyll fluorescence in red. All scale bars are 20 μm. **h** model showing hypothesized mechanism behind vegetative cell leakage or lysis-based calcite precipitation. **i** model showing hypothesized mechanism behind heterocyst contact mediated calcite growth.

In a second instance, shown in Fig. 2c and Supplementary Movie S5, a filament is stressed at a bend. A new crystal nucleates at the bend and grows at the cellular junction. *Anabaena* cells are joined internally within a filament. Therefore, it is likely that the mechanical stress at the bending site allowed cell material to leak out of the cell. These CaCO_3_ precipitation events are likely caused by mechanical confinement or stress to cells, resulting in the leak or entire release of their contents into the surrounding media. Since cyanobacterial cells sequester carbon as carbonate ions, the release of cellular metabolites causes crystal growth. Interestingly, we found that precipitation required these mechanical interactions and was not caused simply by proximity to growing vegetative cells. Fig. 2d and Supplementary Movie S6 shows examples of vegetative cells growing close to or against a seed crystal. The crystal does not increase in size despite ∼26 hours of cellular growth. This result contrasts with previously reported suggestions that cyanobacteria cause MICP due to photosynthetic processes alone^26^.

### Heterocyst contact can result in calcium carbonate crystal growth

A more unique interaction was observed when a seed crystal came into physical contact with a heterocyst cell. A representative example is shown in Fig. 2f and Supplementary Movie S7. Here, a vegetative cell differentiates into a heterocyst, which then physically contacts a seed crystal causing rapid growth. As shown in Fig. 2d, this crystal growth does not occur through contact with a vegetative cell alone. To our knowledge, this observation of heterocyst-mediated MICP has not been previously observed.

To better characterize these events, we visually identified cases where a heterocyst cell contacts a seed crystal. In two out of the seven datasets collected, we clearly observed at least 16 instances when at least one heterocyst contacted a seed crystal. Only instances where we could clearly observe both heterocyst and seed crystal were included in this count. Cases where there was ambiguity, for instance if there were overlapping cells, if the seed crystals or cells were out of the plane of focus, or if a bubble washed out the cells, were ignored. We did not observe heterocyst contact in three replicate movies. However, we note that whether a heterocyst contacts a seed crystal occurs entirely by chance.

While the exact mechanism is currently unknown, whether heterocyst contact causes a crystal to grow appears to depend on when heterocyst contact was made. From these, nine events did not show crystal growth (as can be seen in Supplementary Movie S2), while the remainder did. The contact events which did not result in crystal growth occurred in the early portion of the videos, between 8 and 51 hours. The remaining examples, occurring 37 hours or later, resulted in crystal growth.

### Biotic Calcium Carbonate Crystals are Calcite

To further characterize the crystals, we grew and filmed cells using a Raman-capable microscope. A zoomed in region from the final fame of the full movie (Supplementary Movie S8) is shown in Fig. 3a. The sample was manually registered to identify the crystals tracked in the movie. An area Raman scan was then performed, as shown in Fig. 3b. The images are shown overlaid in Fig. 3c to show overlap of crystal location from Raman microscope data with the end of live growth experiment.

**Fig. 3.**
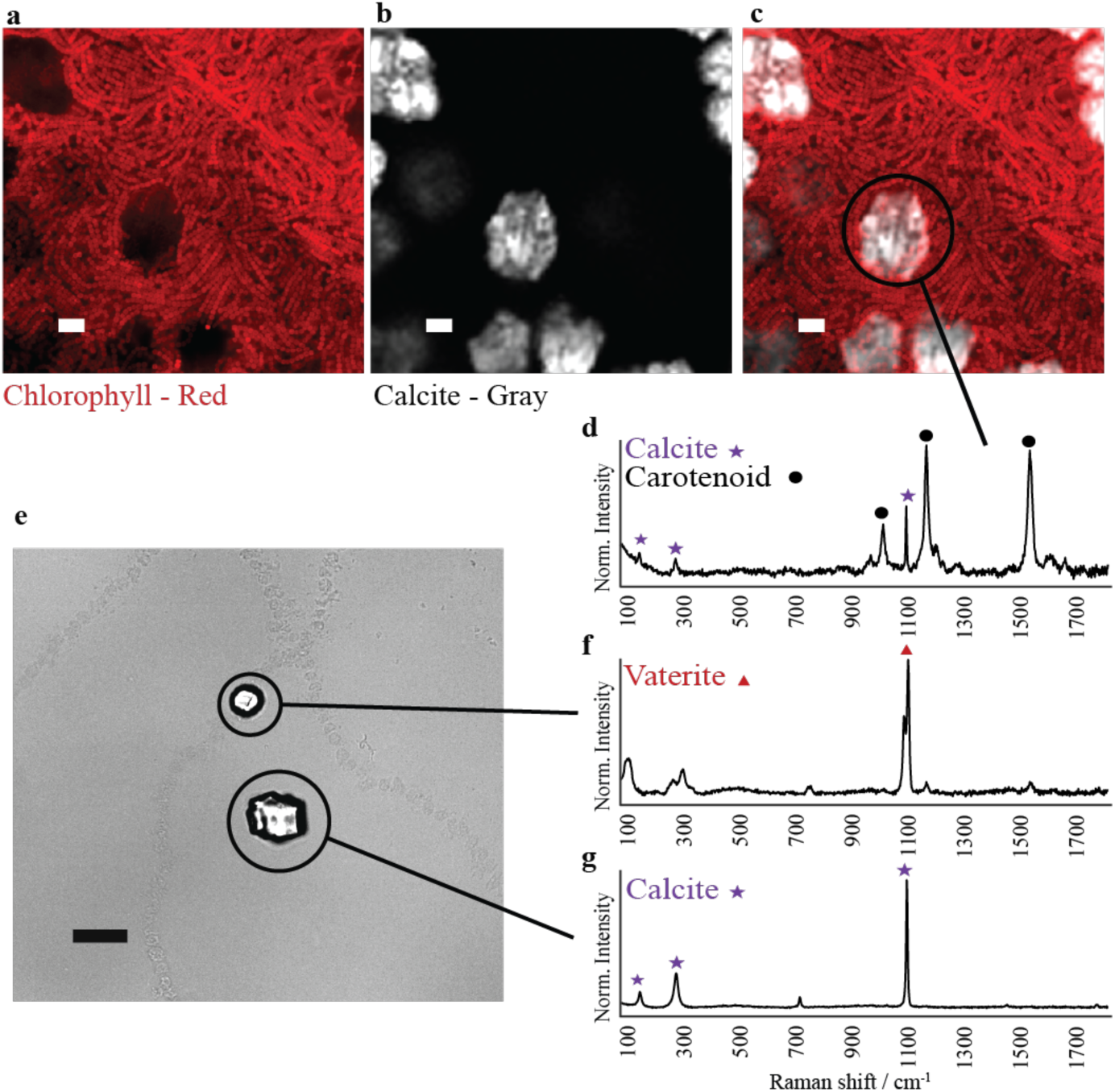
Correlated live-cell and Raman imaging of calcite precipitation by *Anabaena* at single-crystal and single-cell resolution. a-c Area Raman scan with calcite intensity in gray and chlorophyl fluorescence from final frame of live growth experiment in red and an overlay of the two channels. Scale bar is 20 μm. **d** Raman scan of this crystal showing calcite and carotenoid presence. **e** Image of crystals in final frame of time lapse experiment, and **f,g** two corresponding Raman scans of crystals distant from cellular growth showing calcite and vaterite with no carotenoid presence. All scale bars are 20 μm.

Based on the Raman spectra, we found that the crystals consisted primarily of calcite. Interestingly, the crystals also showed traces of carotenoids (Fig. 3d, indicating the presence of cell debris^27^. In comparison, abiotic crystals from a different region in the same sample but without cell proximity were much smaller, and appeared to be composed of either calcite or vaterite (Figs. 2d, 2f, and 2g)^28,29^. Additional Raman scans are shown in Supplementary Fig. 1.

We note observed that in other locations on the same sample without cell growth (e.g., due to filament death as shown in Supplementary Movie S9), no crystal growth was observed beyond the initial abiotic crystal growth as previously mentioned as shown in Fig. 3e. The Raman signature for these crystals (Fig. 3e and f) also appeared to be a mix of vaterite and calcite Fig. 3e,f, which is indicative of CaCO_3_ formation in abiotic environments^30^.

### Calcium carbonate crystals anchor cells during live microscopy experiments

While cyanobacteria are known to cause MICP, its evolutionary benefit is still somewhat unclear. Several speculative hypothesis have been proposed, including as a buffer against the pH rise due to photosynthetic activity or as protection against predation^31–33^. Here, we propose a unique hypothesis that MICP could play a role in anchoring cells growing in tidal regions.

In our datasets, we sometimes observe that a gas bubble will grow across the field of view, particularly in videos where cells have been growing for 12 or more hours. These gas bubbles grow because of oxygen gas release during photosynthesis. The bubbles push cells around in a movie and often wash them out of the field of view, shown in Fig. 4a and Supplementary Movie S10. In Figs. 4b and 4c (and corresponding Supplemental Movies S11 and S12), we observed cases where the presence of calcium carbonate crystals appears to allow cells to anchor themselves against movement.

**Fig. 4.**
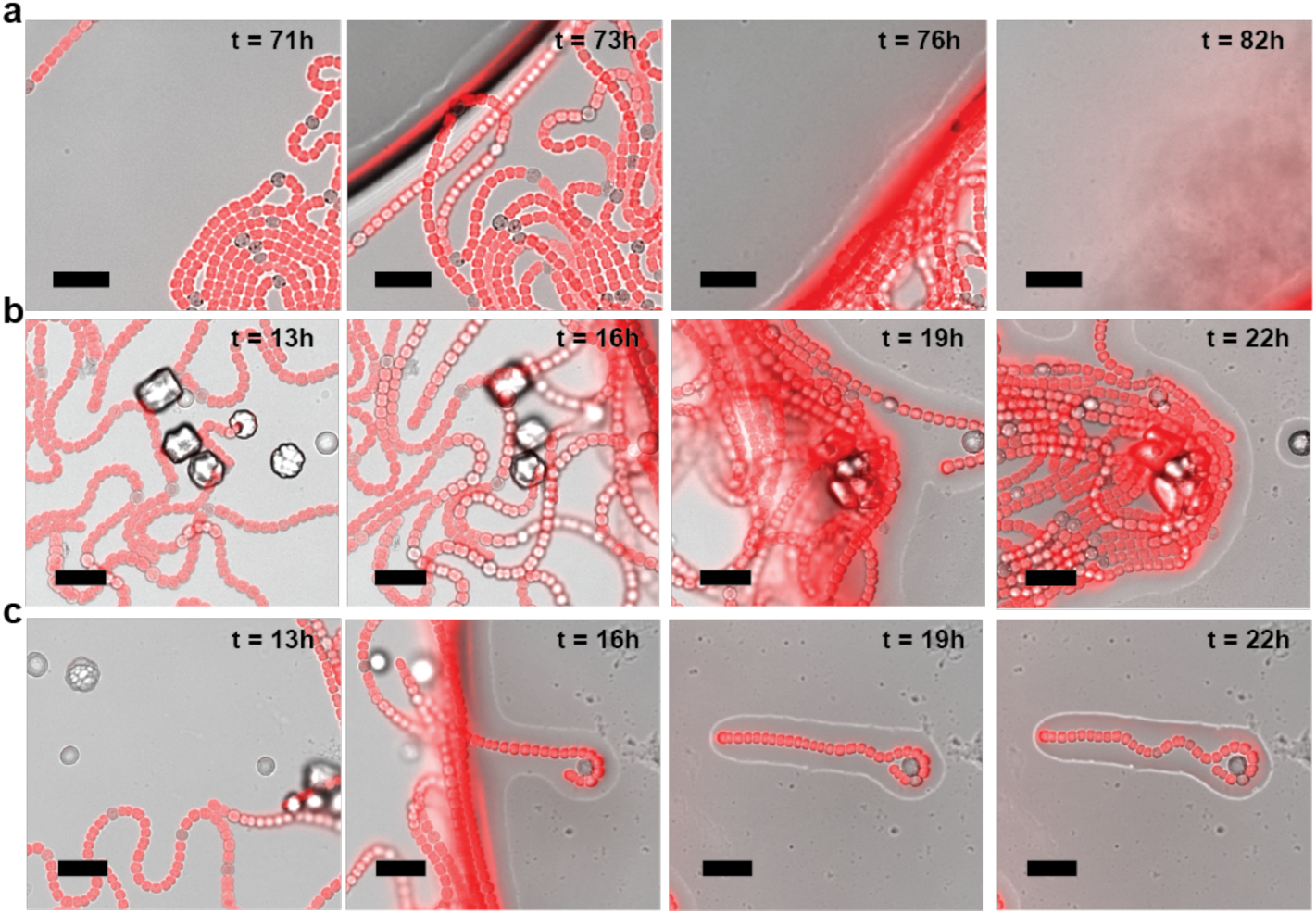
Calcite precipitation acts as an anchoring mechanism against oxygen bubble-induced cell washout. **a** Time-lapse images showing cells being washed away by an oxygen bubble in non-calcium carbonate precipitating conditions. **b, c** Time lapse images showing a bubble passing over cells in calcium carbonate precipitating conditions All scale bars are 20 μm.

## Discussion

A recent study of calcium precipitation by a different strain of *Anabaena* (*A. variabilis*) concluded that MICP could be caused by two distinct mechanisms: (1) *A. variabilis* could produce a precipitating alkaline environment within 24 hours due photosynthetic activity, and (2) *A. variabilis* showed high capacity for adsorbing calcium ions, which could turn the cells into a nucleation agent for crystal formation.^24^ While it is possible that the differences are unique to the species of *Anabaena* studied here, we believe that our results provide greater insight into MICP by these cells.

By monitoring the size of individual crystals, we found that there was no appreciable CaCO_3_ precipitation caused solely by photosynthesis. Instead, MICP appeared to be caused by cell lysis as was previously known, or, more uniquely, by leakage of cellular contents at the junction of mechanically stressed *Anabaena* cells. This stress appears to occur naturally due to bending of the filament and is likely influenced by both cell growth and surface friction of the growth medium^25^. Careful modulation of these conditions could allow MICP to be induced in a controllable manner and should be a target of future studies.

Additionally, we found that MICP was induced by proximity or contact of a heterocyst cell with an existing crystal seed. Interestingly, MICP did not appear to be caused by contact with vegetative cells, which suggests that calcium ion adsorption on the cell surface might not play a key role here, or that calcium might be preferentially bound to heterocysts. Additionally, no preferential calcium carbonate nucleation was observed at the surface of heterocysts, suggesting against calcium ion adsorption as the primary mechanism of heterocyst contact mediated MICP. Alternatively, we hypothesize that MICP could occur due to the nitrogen-fixing activity of the heterocysts. These cells rapidly catalyze the fixation of nitrogen gas into ammonia^34^: *N*_2_ + 16*ATP* + 8*e*^-^ + 8*H*^+^ ⇌ 2*NH*_3_ + *H*_2_ + 16*ADP* + 16*P*_i_. The resulting ammonia is then converted into ammonium ions, 2*NH*_3_ + 2*H*^+^ ⇌ 2*NH*_4_^+^. Thus, 10 protons are consumed per molecule of nitrogen fixed into two molecules of ammonium, which could cause local proton consumption at the cell surface. Further work is required to determine the limits of this mechanism, as it likely works in tandem with environmental factors and is dependent on media conditions.

To our knowledge, MICP caused by heterocyst contact has not been previously observed. This contact-mediated MICP could have evolutionary benefits as an anchoring mechanism. Calcium carbonate is abundant in coastal sediments, forming much of the sand and gravel deposits on beaches^35^. The level of calcium used in these experiments are within reason to those that can be found in these environments,^36^ and at or below that used in most biocementation applications^37,38^. Thus, anchoring could provide a filament with the means to maintain optimal growing conditions by avoiding being washed out by tides. The filamentous nature of *Anabaena* would mean that while a crystallization event would likely result in the death of some cells, this could still be overall beneficial for the rest of the filament.

Overall, obtaining a better understanding of MICP processes in cyanobacteria is important in the search for greener concrete alternatives. Currently, most cement manufacturing involves baking limestone and other minerals in a kiln. Some estimates suggest that cement production contributes ∼5% of the total fossil fuel CO_2_ emissions globally and is a major contributor of global warming^39^. Conversely, biocement manufacturing uses MICP-causing bacteria to precipitate calcium carbonate. However, current biocement manufacturing typically utilize bacteria such as *Sporosarcina pasteurii*, which cause MICP through urea hydrolysis^40^. This process requires fixed nitrogen in the form of urea to be supplied. Urea production is itself a major contributor to global warming, with estimates suggesting that the industry consumes up to 1% of the global energy production and accounts for ∼1% of the global annual CO_2_ emissions as of 2010^41^, and the byproducts of urea hydrolysis releases ammonia and ammonium pollutants into the environment which can adversely affect human and environmental health^42^. By using *Anabaena*, MICP can occur without the need for addition of urea or other ammonia derivatives, as the cells are able to fix atmospheric nitrogen. This represents significant resource savings over alternative biocementation approaches, including those involving other species of cyanobacteria. By providing a better understanding of how MICP is caused by this species, this work could lead to more optimized production yields, reduced resource costs, and decreased harmful byproducts during manufacturing.

## Methods

### Strain and culturing

*Anabaena* sp. ATCC 33047 was grown at 37 °C in BG11 media. All preculturing occurring in 25 mL liquid cultures with 100 rpm orbital shaking in 125 ml baffled flasks with foam stoppers. Liquid and agar cultures were grown in an AL-41 L4 Environmental Chamber (Percival Scientific, Perry, IA) at 37 °C under continuous illumination at 150 μmol photons m^−2^ s^−1^ from cool white fluorescent lamps in air (0.04% CO_2_).

### Time-lapse microscopy

All long-term observation images and videos were obtained using a Nikon TiE inverted wide-field microscope with Perfect Focus System, controlled using NIS Elements AR software (version 5.11.00; 64-bits) with Jobs package. Images were acquired using a digital sCMOS camera (Hamamatsu ORCA-Flash4.0 V2+) with a 20× air objective (Nikon CFI Plan Apochromat Lambda D). Temperature during cell growth in all images was maintained at 37 °C using an Okolab cage Incubator (Okolab). Growth light and trans-illuminating imaging light were supplied from a light-emitting diode (LED) light source (LIDA Light Engine, Lumencor, Beaverton, OR). Epifluorescence imaging light was supplied from a custom-filtered LED light source (Spectra X Light Engine, Lumencor, Beaverton, OR) and delivery was controlled using a synchronized hardware-triggered shutter. Cells for imaging were grown in liquid culture to ∼1.00 OD at 730 nm. Precultures were grown in BG11 media. 25 μL of 1 M CaCl_2_ was added to the sample side of a 1% agarose imaging pad (1 mL) BG11_-N-Buffer_ (BG11 media without fixed nitrogen or buffer, pH 7.8) and allowed to fully dry/dissolve (for non-MICP condition movies, the imaging pad was made with BG11_-N_ (BG11 media without fixed nitrogen) media and no CaCl_2_ or sodium bicarbonate were added). Three to five 2 μL drops of cells were added to the imaging side of the pad and allowed to dry. The pad was flipped into the imaging dish, and an additional 25 μL of 1 M sodium bicarbonate was added to the top of the pad. The imaging dish (Ibidi μ-dish 35mm glass) was immediately sealed with parafilm, and placed into the microscope environmental chamber at 37 °C. Cells were grown under 37 °C and 150 μmol photons m^−2^ s^−1^ 640 nm light; this light was on at all times except during fluorescence imaging.

All time lapse imaging performed on Nikon Plan Apo (wavelength) 20×. Brightfield illuminated by Lida at ExW: 450 nm, power 2.0, ExW: 550 nm, power 2.0, ExW: 640 nm, power 2.0. Cy5 excited at 640 nm, power 5.0. When present, CFP excited at 440 nm, power 50.0. RFP excited at 555 nm, power 10.0. Frames were captured either every 20 or 30 minutes. A total of seven experimental replicates were acquired over the course of two years (see Supplementary Table S1).

### Correlated live-cell and Raman Microscopy

To obtain the correlated light and Raman microscopy images shown in Fig. 3, cells were initially filmed using a light microscope, as described above. A large image was taken at the end of the time course. The sample was then transferred to the Raman microscope (Horiba LabRAM HR Evolution confocal Raman spectrometer with 532/785 nm lasers). Scans of the Raman spectrum across the sample were with either a 10× or 50x objective. Raman scans were performed using a 532 nm laser and scanning from 84 to 1786 cm^-1^. All Raman scans were performed the same day as live growth microscopy and samples were stored in the dark during transfer. Additional Raman scans shown in Supplementary Fig. S1. The resulting Raman scan was then manually correlated with the light microscopy image to identify the mineral identity of each crystal. Raman scans were processed with the Horiba Labspec 6 (Version 6.7.1.10 64-bit) software using a 6-degree iterative polynomial fit. Raman scan was analyzed using a least squared fit using n-member regions within the spectra. The resulting Raman map was colored based on pixel scan similarity to known calcite spectra.

### Image analysis

Image segmentation was performed using a modified version of CyAN^43^ that has been further customized for filamentous cyanobacteria. Filament analysis and measurements performed using MATLAB scripts based on functions present within CyAN and made available online at https://github.com/ChrisBrininger/Cyanobacteria. All computational analysis was performed with MATLAB version R2020b.

## Acknowledgements

The authors would like to dedicate this publication to the memory of Prof. Jeffrey Cameron, who was the principal investigator through most of this study. Jeffrey Cameron was a passionate scientist and incredible mentor, and his guidance on this and many other projects made these discoveries possible. We also thank Dr. Hamadri Pakrasi for the *Anabaena* sp. ATCC 33047 strain and to Eric Ellison for training and correspondence on microscope use and data treatment. This work was supported in part by the NIH Biophysics Training Grant T32 GM145437, and the Department of Energy under grants DOE DE-SC0018368 and DE-SC0020361 to J.C.C. Raman spectroscopic analyses were performed at the Raman Microspectroscopy Lab, Department of Geological Sciences, University of Colorado Boulder (RRID: SCR_019305).

## Author contributions

J.C.C., C.M.B., and E.E. conceptualized the study. J.C.C. and C.M.B. designed the experiments. C.M.B. and E.B.J. ran and acquired the fluorescence microscopy data. C.M.B. performed the Raman microscopy experiments. C.M.B and J.W.T. analyzed the resulting imaging datasets. C.M.B., J.W.T., and J.C.C. wrote the initial manuscript. C.M.B, J.W.T., E.J.B., and E.E. revised and approved the final manuscript.

## Competing Interests

J.C.C. was a co-founder and held equity in Prometheus Materials Inc.

### Data availability

Data and materials are available upon request to CMB or JWT.

