## Supplementary Figures for "Calcite Precipitation by a Nitrogen-Fixing Cyanobacterium"

### Supplementary Information

**Supplementary Video S1**

**Time-lapse imaging of cell growth of *Anabaena* growing without supplemental nitrogen ($BG11_{-N}$).** Images obtained using a 20x objective. The movie shows cells growing in standard growth conditions. The movie is merged images of brightfield in gray and chlorophyll in red.

**Supplementary Video S2**

**Time-lapse imaging of cell growth of *Anabaena* performing MICP.** *Anabaena* grown without supplemental nitrogen and with supplemented calcium carbonate precipitating conditions ($BG11_{-N-Buffer}$+Ca) on a 20x objective. The movie shows cells growing and one calcium carbonate crystal growing in response. The movie is merged images of brightfield in gray and chlorophyll in red.

**Supplementary Video S3**

**Wide field of view time-lapse imaging of cell growth of *Anabaena* performing MICP.** *Anabaena* grown without supplemental nitrogen and with supplemented calcium carbonate precipitating conditions ($BG11_{-N-Buffer}$+Ca) on a 20x objective. The movie shows the movie from which the crystal masks and growth traces were generated. The movie is merged images of brightfield in gray and chlorophyll in red.

**Supplementary Video S4**

**Close up view time-lapse imaging of cell growth of *Anabaena* performing MICP through vegetative cell mechanical lysis due to cell entrapment.** *Anabaena* grown without supplemental nitrogen and with supplemented calcium carbonate precipitating conditions ($BG11_{-N-Buffer}$+Ca) on a 20x objective. The movie shows cells growing and a vegetative cell lysis mediated crystal growth event. The movie is merged images of brightfield in gray and chlorophyll in red.

**Supplementary Video S5**

**Close up view time-lapse imaging of cell growth of *Anabaena* performing MICP through vegetative cell leakage at a filament bend.** *Anabaena* grown without supplemental nitrogen and with supplemented calcium carbonate precipitating conditions ($BG11_{-N-Buffer}$+Ca) on a 20x objective. The movie shows cells growing and the spontaneous nucleation and growth of a crystal from what appears to be a vegetative cell leakage event. The movie is merged images of brightfield in gray and chlorophyll in red.

**Supplementary Video S6**

**Close up view time-lapse imaging of cell growth of *Anabaena* vegetative cell contacting a calcium carbonate crystal without crystal growth.** *Anabaena* grown without supplemental nitrogen and with supplemented calcium carbonate precipitating conditions ($BG11_{\text{-N-Buffer}}$+Ca) on a 20x objective. The movie shows cells growing and no corresponding crystal growth upon contact between a crystal and a vegetative cell. The movie is merged images of brightfield in gray and chlorophyll in red.

**Supplementary Video S7**

**Close up view time-lapse imaging of cell growth of *Anabaena* performing MICP through heterocyst contact with a crystal seed.** *Anabaena* grown without supplemental nitrogen and with supplemented calcium carbonate precipitating conditions ($BG11_{\text{-N-Buffer}}$+Ca) on a 20x objective. The movie shows cells growing and a heterocyst coming in contact with a calcium carbonate crystal, resulting in further crystal growth. The movie is merged images of brightfield in gray and chlorophyll in red.

**Supplementary Video S8**

**Time-lapse imaging of cell growth of *Anabaena* MICP at the location used for Raman scanning.** *Anabaena* grown without supplemental nitrogen and with supplemented calcium carbonate precipitating conditions ($BG11_{\text{-N-Buffer}}$+Ca) on a 20x objective. The movie shows cells growing at high density and resulting in many crystal growth events, which were later scanned with Raman microscopy. The movie is merged images of brightfield in gray and chlorophyll in red.

**Supplementary Video S9**

**Time-lapse imaging of an *Anabaena* filament which dies and does not result in MICP.** *Anabaena* grown without supplemental nitrogen and with supplemented calcium carbonate precipitating conditions ($BG11_{-N-Buffer}$+Ca) on a 20x objective. The movie shows cells which do not grow, and there is no resulting crystal growth. The movie is merged images of brightfield in gray and chlorophyll in red.

**Supplementary Video S10**

**Time-lapse imaging of *Anabaena,* growing without supplemental MICP conditions, being washed away by a bubble.** *Anabaena* grown without supplemental nitrogen ($BG11_{-N}$) on a 20x objective. on a 20x objective. The movie shows cells growing until an oxygen bubble passes over the field of view, which then pushes all cells out of focus and/or view. The movie is merged images of brightfield in gray and chlorophyll in red.

**Supplementary Video S11**

**Time-lapse imaging of an *Anabaena* growing with supplemental MICP conditions, performing MICP and avoiding being washed away by a bubble.** *Anabaena* grown without supplemental nitrogen and with supplemented calcium carbonate precipitating conditions ($BG11_{-N-Buffer}$+Ca) on a 20x objective. The movie shows cells growing until an oxygen bubble passes over the field of view, at which point cells are anchored from movement by crystal presence/ growth. The movie is merged images of brightfield in gray and chlorophyll in red.

**Supplementary Video S12**

**Time-lapse imaging of an *Anabaena* growing with supplemental MICP conditions, performing MICP at the point of contact between a heterocyst and a crystal seed, and avoiding being washed away by a bubble.** *Anabaena* grown without supplemental nitrogen and with supplemented calcium carbonate precipitating conditions ($BG11_{-N-Buffer}$+Ca) on a 20x objective. The movie shows cells growing until and oxygen bubble passes over the field of view, at which point cells are anchored from movement by crystal presence/ growth, with one specific crystal focused on which grows due to heterocyst contact mediated calcite growth. The movie is merged with images of brightfield in gray and chlorophyll in red.

**Supplementary Table 1. List of experimental replicate datasets, acquisition date, number of separate fields of view within the replicate movie, and total duration recorded of each movie.** Each movie was recorded with 30 minutes between frames, except for replicate 7, which was acquired at 20 minutes between frames.

| Replicate # | Date | Number of Fields of View | Duration |
|---|---|---|---|
| 1 | 03/01/2021 | 2 | 95.5 Hours |
| 2 | 04/18/2021 | 3 | 50.5 Hours |
| 3 | 05/31/2021 | 5 | 161.5 Hours |
| 4 | 04/11/2022 | 3 | 100.5 Hours |
| 5 | 04/20/2022 | 4 | 114.5 Hours |
| 6 | 06/10/2022 | 4 | 94 Hours |
| 7 | 07/19/2023 | 5 | 117.7 Hours |

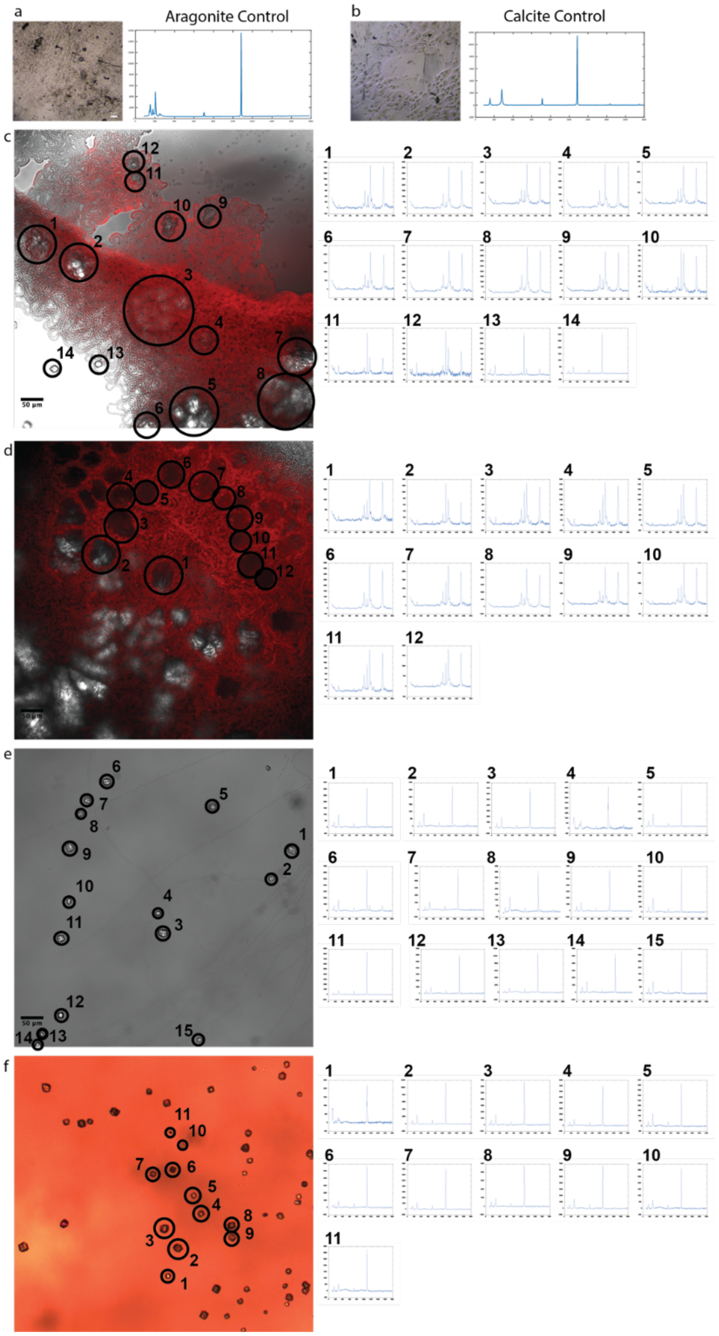


**Supplementary Figure S1. Raman microscopy scans of calcium carbonate precipitated with and without cell proximity.** Images taken using a 20x objective, and scans taken using 50x objective at the center of circled crystals. **a, b** Raman scans of aragonite and calcite control. **c-d** Raman scans of targeted crystals from live growth microscopy experiment. Images taken using a 20x objective, and scans taken using 50x objective at the center of circled crystals. **f** Raman scans from distal region of same growth pad used in live growth microscopy experiment shown in c-d, but with no cells within view. Image taken using a 10x objective, and scans taken using 50x objective at the center of the circled crystals.
